# C9orf72 Repeat Expansion Induces Metabolic Dysfunction in Human iPSC-Derived Microglia and Modulates Glial–Neuronal Crosstalk

**DOI:** 10.1101/2025.03.14.643263

**Authors:** Marika Mearelli, Insa Hirschberg, Francesca Provenzano, Christin Weissleder, Mafalda Rizzuti, Linda Ottoboni, Stefania Corti, Michela Deleidi

## Abstract

The C9orf72 hexanucleotide repeat expansion mutation is the most common genetic cause of amyotrophic lateral sclerosis (ALS) and frontotemporal dementia, but its cell type-specific effects on energy metabolism and immune pathways remain poorly understood. Using induced pluripotent stem cell (iPSC)-derived motor neurons, astrocytes and microglia from C9orf72 patients and their isogenic controls, we investigated metabolic changes at the single-cell level under basal and inflammatory conditions. Our results showed that microglia are particularly susceptible to metabolic disturbances. While C9orf72 motor neurons exhibited impaired mitochondrial respiration and reduced ATP production, C9orf72 microglia presented pronounced increases in glycolytic activity and oxidative stress, accompanied by the upregulation of the expression of key metabolic enzymes. These metabolic changes in microglia were exacerbated by inflammatory stimuli. To investigate how these changes affect the broader cellular environment, we developed a human iPSC-derived triculture system comprising motor neurons, astrocytes and microglia. This model revealed increased metabolic activity in all cell types and highlighted that microglia-driven metabolic reprogramming in astrocytes contributes to the vulnerability of motor neurons under inflammatory conditions. Our findings highlight the central role of microglia in driving metabolic dysregulation and intercellular crosstalk in ALS pathogenesis and suggest that targeting metabolic pathways in immune cells may provide new therapeutic avenues.

## Introduction

Amyotrophic lateral sclerosis (ALS) is the most common adult-onset motor neuron disease (MND) and is characterized by the progressive degeneration and death of motor neurons (MNs) in the cerebral cortex, brainstem, and spinal cord. ALS pathogenesis is multifactorial and involves diverse mechanisms that contribute to the selective vulnerability and loss of MNs. Traditionally considered a disease driven primarily by the intrinsic susceptibility of MNs to autonomous cell death, increasing evidence highlights the importance of noncell-autonomous mechanisms in MN degeneration [1]. In particular, interactions among neurons, astrocytes, and immune cells have emerged as pivotal contributors to disease onset and progression [2]. ALS is characterized by widespread neuroinflammation, including astrogliosis, microglial activation, and peripheral immune cell infiltration at sites of neuronal degeneration. Notably, autoimmune diseases often precede the onset of ALS/frontotemporal dementia (FTD) [3, 4], and microglial activation at disease onset is correlated with the rate of motor decline [5, 6].

Although most ALS cases are sporadic, several gene mutations are associated with familial forms of the disease. Among these, the expansion of a noncoding hexanucleotide repeat (GGGGCC) in the *C9orf72* gene is the most common genetic cause of ALS and FTD [7, 8]; this mutation accounts for 20–40% of familial ALS cases and 2–8% of sporadic cases, with a higher prevalence among individuals of European descent [9]. The high frequency of *C9orf72* mutations in both ALS and FTD has stimulated extensive research into its role in neuronal damage and degeneration. A critical question is whether and how C9orf72 contributes to the risk and progression of the disease. *C9orf72* is ubiquitously expressed, including in peripheral myeloid cells and microglia [10]. The loss of C9orf72 in mice enhances proinflammatory responses in peripheral myeloid cells and microglia [11, 12]. Emerging evidence also highlights the critical role of C9orf72 in intercellular communication and metabolic regulation, with C9orf72 being a key regulator of mitochondrial function and cellular energy homeostasis [13]. Interestingly, compared with control astrocytes, ALS-induced astrocytes display a dysregulated metabolic profile characterized by a significant loss of metabolic flexibility [14]. Additionally, C9orf72 deficiency has been associated with impairments in autophagy and lysosomal function—processes essential for cellular homeostasis and the clearance of pathogenic aggregates capable of propagating between cells [15]. The C9orf72-mediated regulation of inflammatory signaling in macrophages and microglia may influence the local metabolic microenvironment, potentially disrupting the neuron‒glia interactions critical for disease progression. Such disruptions may exacerbate energy deficits in MNs, further increasing their susceptibility to degeneration. Understanding how C9orf72 affects cell type-specific metabolism and intercellular communication is essential for identifying novel therapeutic strategies for ALS and related diseases.

To address these issues, we generated C9orf72 mutant human induced pluripotent stem cell (iPSC)-derived MNs and glial cells (microglia and astrocytes) in both monoculture and coculture systems. We used spectral flow cytometry for high-dimensional single-cell analysis, integrating metabolic flow cytometry (Met-Flow) to comprehensively assess cell type-specific cellular metabolism. This study provides an in-depth metabolic analysis of human C9orf72 mutant iPSC-derived MNs and glia, providing novel insights into C9orf72-related metabolic dysregulation, neuron‒glia interactions and their contributions to disease pathogenesis.

## Results

### The C9orf72 repeat expansion mutation exerts a cell type-specific effect on energy metabolism in human iPSC-derived motor neurons, astrocytes, and microglia

To investigate the cell type-specific impact of the *C9orf72* repeat expansion mutation on energy metabolism, we used three independent C9orf72 patient iPSC lines (BS6, C9-1; CS29iALS, C9-2; and CS52iALS, C9-3) and their corresponding isogenic controls (BS6 2H9, Iso-1; CS29iALS-Iso, Iso-2; and CS52iALS-Iso, Iso-3). The iPSCs were differentiated into highly enriched spinal cord MNs, microglia, and astrocytes. The C9orf72 repeat expansion mutation did not alter the differentiation potential of iPSCs (Figures 1-3). Next, we analyzed the oxygen consumption rate (OCR) in C9orf72 MNs, microglia, and astrocytes using Seahorse with the sequential addition of the complex V/ATP synthase inhibitor oligomycin, the protonophore CCCP, and the complex III inhibitor antimycin A to measure several parameters of mitochondrial function. We detected a significant reduction in maximal mitochondrial respiration in C9orf72 MNs compared with isogenic controls (Figure 1B). In parallel, ATP levels were significantly lower in C9orf72 MNs than in isogenic controls (Figure 1B, Supplementary Figure 1). Compared with isogenic controls, C9orf72-derived astrocytes and microglia presented nonsignificant differences in the OCR, with a trend toward increased respiration (Figure 2B, 3B). ATP levels were significantly increased in C9orf72-derived astrocytes and microglia (Supplementary Figure 1B). Next, we assessed the glycolytic rate in iPSC-derived MNs, astrocytes, and microglia. While MNs did not exhibit a significant change, C9orf72-derived microglia presented a greater extracellular acidification rate (ECAR) than isogenic controls did, whereas no significant increase was observed for C9orf72-derived astrocytes (Supplementary Figure 1B).

**Figure 1.**
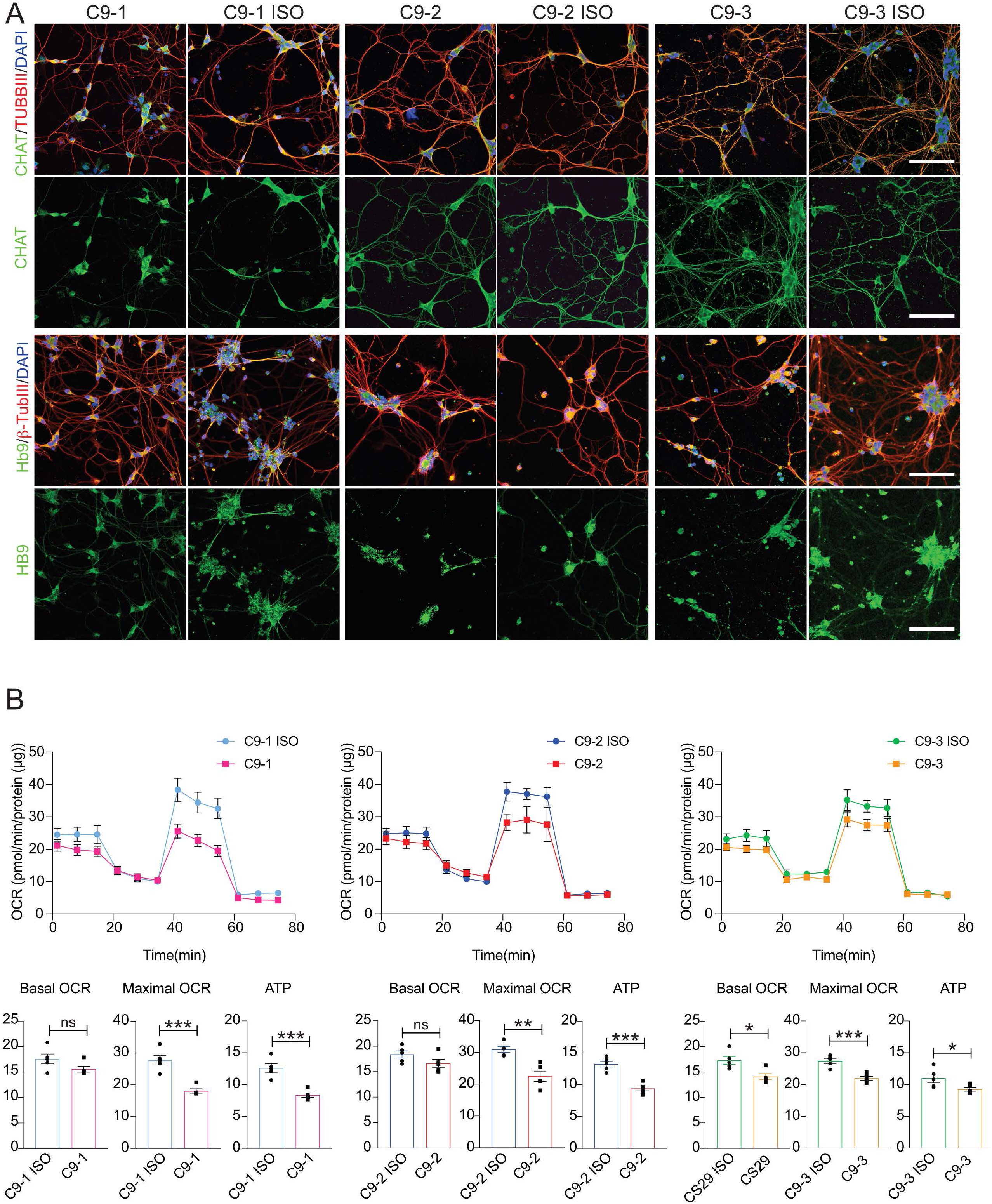
*C9orf72* expansion reduces mitochondrial respiration in motor neurons (MNs) derived from iPSCs. **A)** Representative confocal images showing the immunocytochemical staining of CHAT and HB9 (green) in C9orf72 (C9) and paired isogenic control (C9-ISO) iPSC-derived MNs; TUBB3 (red) and nuclei (DAPI; blue). Scale bar, 100 µm. **B)** Mitochondrial respiration in isogenic C9 and C9-ISO MNs. Oxygen consumption rates (OCRs) were measured over time after the sequential addition of oligomycin, CCCP and rotenone/antimycin A. Values for the basal OCR, ATP-linked OCR, and maximal OCR are expressed as pmol/min/protein (µg). Mean ± SEM; unpaired two-tailed t test; *** P<0.001, ** P<0.01, and * P<0.05; n=5 independent experiments.

**Figure 2.**
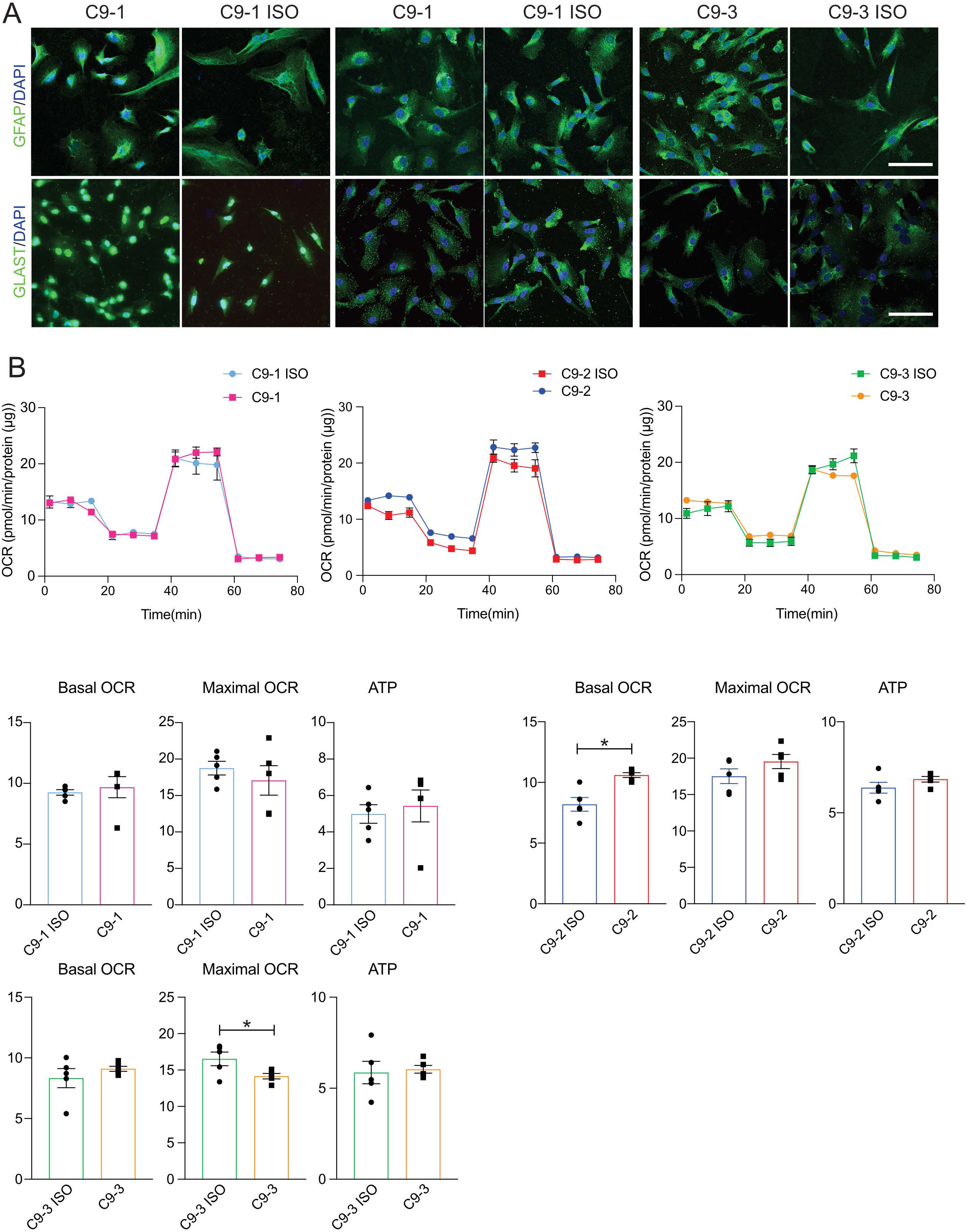
Differentiation and mitochondrial bioenergetic characterization of C9orf72 isogenic iPSC-derived astrocytes. **A)** Representative confocal images showing the immunocytochemical staining of GFAP (green) and GLAST (green) in isogenic C9orf72 (C9) and paired isogenic control (C9-ISO) iPSC-derived astrocytes; nuclei (DAPI; blue). Scale bar, 100 µm. **B)** Mitochondrial respiration in C9 and C9-ISO astrocytes. Oxygen consumption rates (OCRs) were measured over time after the sequential addition of oligomycin, CCCP and rotenone/antimycin A. Values for the basal OCR, ATP-linked OCR, and maximal OCR are expressed as pmol/min/protein (µg). Mean ± SEM; unpaired two-tailed t test; * P<0.05; n=5 independent experiments.

**Figure 3.**
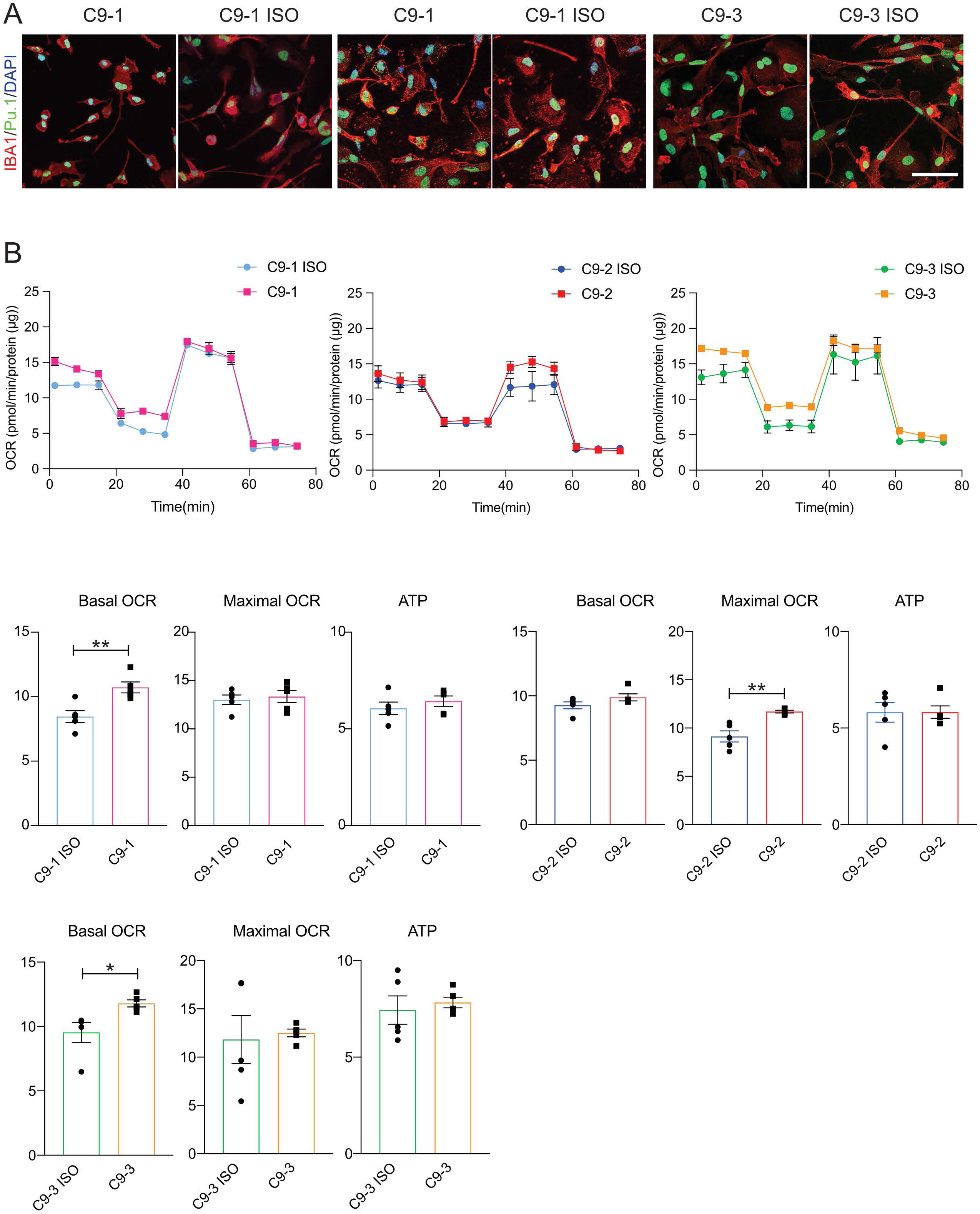
Differentiation and mitochondrial bioenergetic characterization of C9orf72 isogenic iPSC-derived microglia. **A)** Representative confocal images showing the immunocytochemical staining of IBA1 (red) and Pu.1 (green) in C9orf72 (C9) and paired isogenic control (C9-ISO) iPSC-derived microglia; nuclei (DAPI, blue). Scale bar, 100 µm. **B)** Mitochondrial respiration in C9 and C9-ISO microglia. Oxygen consumption rates (OCRs) were measured over time after the sequential addition of oligomycin, CCCP and rotenone/antimycin A. Values for the basal OCR, ATP-linked OCR, and maximal OCR are expressed as pmol/min/protein (µg). Mean ± SEM; unpaired two-tailed t test; ** P<0.01 and * P<0.05; n=5 independent experiments.

### Single-cell metabolic analysis reveals enhanced glycolysis and oxidative stress responses in C9orf72 mutant microglia

To validate the metabolic features and assess the cell type-specific impact of *C9orf72* expansion, we used MET-flow, a single-cell metabolic analysis strategy [16]. We combined a panel of different metabolic enzymes involved in different pathways: glucose uptake (glucose transporter 1, GLUT1), the oxidative pentose phosphate pathway (glucose 6-phosphate dehydrogenase, G6PD), glycolysis (hexokinase-1, HK1), and antioxidant response pathways (peroxiredoxin 2, PRDX2) (Figure 4A). MNs and astrocytes from both C9orf72 and isogenic control cultures showed no significant differences in the expression of GLUT1, HK1, G6PD, or PRDX2, suggesting that neuronal and astrocyte metabolism might be relatively unaffected by *C9orf72* expansion under basal conditions (Figure 4B). In contrast, microglia from C9orf72 cultures presented significant metabolic changes. GLUT1 and PRDX2 expression was significantly higher in C9orf72 microglia than in control microglia, indicating increased glycolytic activity and oxidative stress (Figure 4B, Supplementary Figure 2A). These findings suggest that microglia are particularly susceptible to metabolic disruptions associated with *C9orf72* expansion, even in the absence of external inflammatory stimuli.

**Figure 4.**
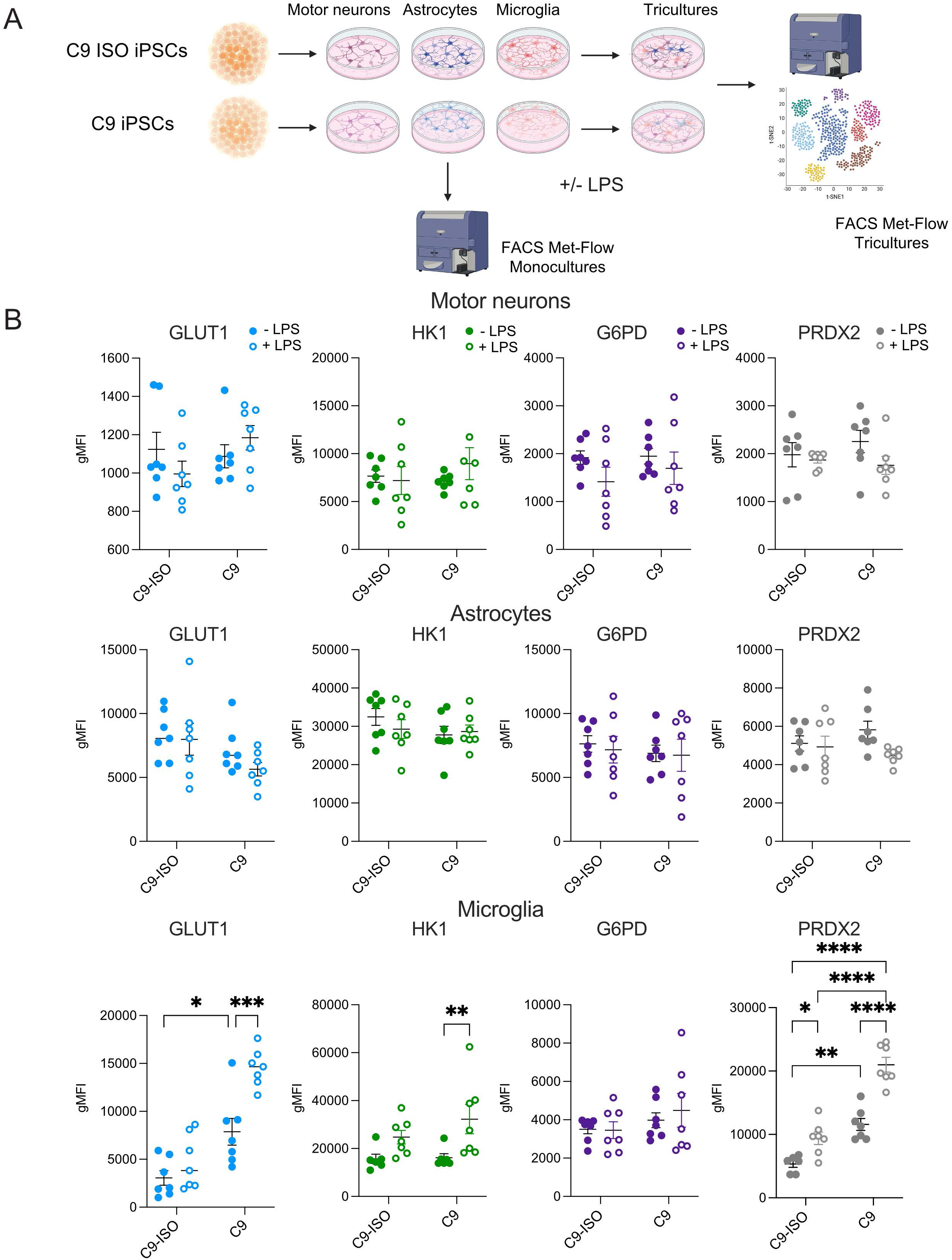
MET-flow-based metabolic analysis revealed divergent metabolic profiles in C9orf72 isogenic iPSC-derived MNs, astrocytes, and microglia in monocultures. **(A)** Schematic of the experimental workflow. C9-ISO iPSCs and C9 iPSCs (C9-1 and C9-2 and corresponding isogenic controls) were differentiated into MNs, astrocytes, and microglia. These cells were then either left untreated or treated with LPS and analyzed as monocultures or as part of a triculture system. Using MET-Flow, we evaluated the expression of key metabolic enzymes— GLUT1, G6PD, HK1, and PRDX2—in MNs, astrocytes, and microglia derived from C9-ISO iPSCs and C9 iPSCs. **(B)** Quantitative analysis of metabolic markers—GLUT1, HK1, G6PD, and PRDX2—in MNs, astrocytes, and microglia derived from C9 and C9 ISO iPSC lines under basal conditions and after LPS treatment. The data are expressed as the geometric mean fluorescence intensity (gMFI). Mean ± SEM; two-way ANOVA with Bonferroni correction; ***P < 0.001, **P < 0.01, and *P < 0.05; n=6-7 independent experiments.

### Single-cell metabolic analysis reveals a role for C9orf72 in microglial metabolic reprogramming under inflammatory conditions

To characterize changes in cellular metabolism under inflammatory conditions, we performed MET-flow analysis by quantifying rate-limiting metabolic enzymes after LPS stimulation. Compared with the isogenic controls, in C9orf72 microglia, LPS treatment significantly increased the expression of GLUT1, HK1 and PRDX2, indicating an enhanced metabolic and oxidative stress response (Figure 4B, Supplementary Figure 2B). C9orf72 MNs and astrocytes did not significantly differ in GLUT1, HK1, G6PD or PRDX2 expression after LPS stimulation (Figure 4B). These results suggest that, even though both glial and neuronal cell types express TLR4 [17–19], LPS-induced inflammation drives differential metabolic responses in MNs, astrocytes, and microglia, with the significant upregulation of the expression of key metabolic enzymes in C9orf72 microglia. The increased expression of GLUT1, G6PD and PRDX2 in microglia suggests a potential mechanism for metabolic dysregulation and oxidative stress in response to inflammatory stimuli in neurodegenerative disease processes.

### Neuronal‒glial metabolic interactions in an iPSC-derived triculture system are linked to increased cellular metabolic activity

Neuronal‒glial interactions play a critical role in modulating inflammatory responses in the central nervous system (CNS), and changes in their interactions have been implicated in ALS [20]. To assess the role of C9orf72 in neuronal–glial metabolic interactions, we developed a human iPSC-derived triculture system. This system consists of 35% iPSC-derived MNs, 55% astrocytes, and 5% microglia (Figure 5A). Immunofluorescence imaging confirmed the presence of MNs, astrocytes, and microglia within the triculture system, with comparable cellular distributions between C9orf72 and isogenic triculture conditions (Figure 5A). After 10 days of coculture, the metabolic profiles of MNs, astrocytes, and microglia derived from iPSCs were assessed in tricultures using Met-Flow (Supplementary Figures 3, 4). Compared with monocultures, MNs in tricultures presented significantly greater expression of GLUT1, G6PD, and HK1, indicating a global increase in metabolic activity when these cells are in a multicellular environment (Supplementary Figure 4). In contrast, the PRDX-2 levels were lower in MNs in the triculture system than in those in the monoculture system, suggesting a reduction in oxidative stress (Supplementary Figure 4). In the triculture system, astrocytes showed a significant decrease in GLUT1 expression (Supplementary Figure 4). Those findings suggest that astrocytes in a multicellular context may experience a slight shift in glucose metabolism but not in oxidative stress levels. Similarly, in microglia, HK1 and G6PD expression was significantly higher in tricultures than in monocultures, highlighting the increased metabolic and oxidative activity of these cells when they interact with other cell types, possibly driven by signals from neighboring MNs (Supplementary Figure 4). Collectively, these findings suggest that the cellular microenvironment plays a critical role in modulating the metabolic state of iPSC-derived MNs, astrocytes, and microglia, increasing the metabolic activity of the different cell types studied.

**Figure 5.**
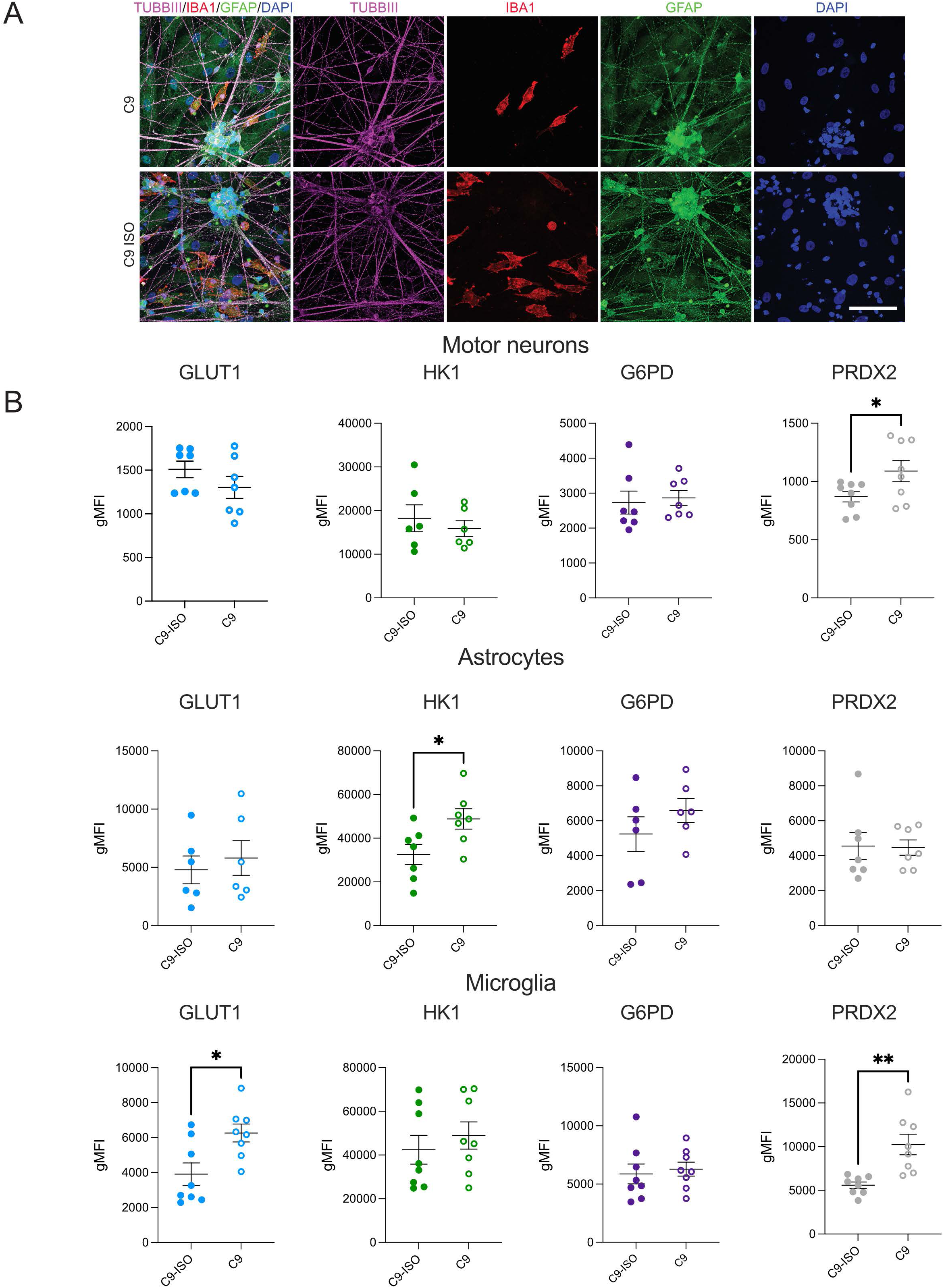
Metabolic profiling of iPSC-derived neuronal–glial tricultures reveals microglial susceptibility to C9orf72-associated disruptions. **(A)** Representative immunofluorescence images showing the cellular composition of C9-ISO and C9 (C9-1 and C9-2 and corresponding isogenic controls) tricultures. The cells were stained for TUBB3 (magenta) to identify MNs; IBA1 (red) to identify microglia; GFAP (green) to identify astrocytes; and DAPI (blue) to identify nuclei. The individual channels for TUBB3, IBA1, and GFAP are shown to the right. Scale bars, 100 µm. **(B)** Met-Flow analysis of metabolic markers in isogenic iPSCs under basal conditions. The metabolic markers GLUT1, HK1, G6PD, and PRDX2 were assessed in MNs, astrocytes, and microglia using Met-Flow. The data are presented as the geometric mean fluorescence intensity (gMFI) for each marker, comparing C9orf72-expanded tricultures (C9) with isogenic controls (C9 ISO Tri). Mean ± SEM; unpaired two-tailed t test; *P < 0.05 and **P < 0.01; n=7 independent experiments.

### The metabolic profiling of iPSC-derived neurons and glia reveals microglial susceptibility to C9orf72-associated metabolic disruptions under basal conditions

To investigate the baseline metabolic profile of iPSC-tricultures with *C9orf72* expansion in the absence of inflammatory stimuli, we utilized Met-Flow to assess the expression levels of key metabolic markers in MNs, astrocytes, and microglia. Under basal conditions, significant metabolic differences were observed in C9orf72-expanded MNs compared with their isogenic controls. In MNs, the expression of PRDX2, an antioxidant enzyme, was significantly increased in C9orf72 cultures, indicating a potential increase in oxidative stress even in the absence of inflammatory stimuli (Figure 5B). No significant changes were detected in the expression of GLUT1, HK1, or G6PD in MNs between C9orf72 and isogenic conditions. Compared with those derived from isogenic controls, astrocytes derived from C9orf72 cultures presented significant increases in HK1 levels, suggesting an alteration in glycolytic metabolism in these cells. However, no significant differences in GLUT1, G6PD, or PRDX2 levels were detected in astrocytes (Figure 5B). Microglia exhibited the most pronounced metabolic changes, with the significant upregulation of both GLUT1 and PRDX2 expression in C9orf72 cultures, indicating an enhanced glycolytic response and heightened oxidative stress. Those results suggest that microglia in C9orf72-expanded cultures may be particularly metabolically vulnerable even in the absence of an external inflammatory trigger (Figure 5B). Overall, our findings highlight the presence of baseline metabolic alterations in iPSC-tricultures with C9orf72 expansion, particularly in microglia and astrocytes, which may contribute to disease pathogenesis even prior to the onset of inflammation.

### C9orf72 microglia drive astrocytic metabolic changes and contribute to motor neuron loss under inflammatory conditions

Next, MET-flow analysis was conducted to examine single-cell metabolic changes in iPSC-tricultures containing C9orf72 expansions compared with those in isogenic controls following LPS treatment. Quantitative metabolic analysis revealed distinct metabolic alterations in astrocytes and microglia harboring C9orf72 expansion compared with isogenic controls. In astrocytes, a significant increase in GLUT1 and HK1 was observed in C9orf72 cells compared with isogenic controls (Figure 6A), suggesting an enhanced glycolytic response under inflammatory conditions; furthermore, the PRDX2 levels were markedly increased, indicating a heightened oxidative stress response (Figure 6A). Similar trends were observed in microglia, where GLUT1, HK1, and PRDX2 levels were significantly elevated in C9orf72 cells, a finding that was consistent with a metabolic shift toward increased glucose metabolism and oxidative stress management (Figure 6A). In contrast, MNs did not exhibit significant metabolic changes in response to LPS treatment, regardless of *C9orf72* expansion. These results suggest that astrocytes and microglia, rather than MNs, may exhibit significant metabolic vulnerabilities in response to inflammatory stress in the context of *C9orf72* expansion, potentially contributing to the neurodegenerative processes observed in related pathologies. The activation of astrocytes is often correlated with the severity of disease phenotypes in mouse models of ALS, suggesting possible microglia‒astrocyte crosstalk that regulates the astrocytic transition from neuroprotective to neurotoxic [21]. Consistent with these findings, our data suggest that activated C9orf72 microglia drive metabolic changes in astrocytes. To explore whether cross-talk between activated microglia and astrocytes has a neurotoxic effect on MNs, we quantified neuronal cell counts in both monoculture and triculture systems. Under basal conditions, MN numbers did not differ between C9orf72 and isogenic controls in either monoculture or the triculture system (Figure 6B-D). However, following LPS treatment in tricultures, we observed a moderate but significant reduction in MN numbers (Figure 6C). Interestingly, treatment of monocultures with conditioned medium obtained from LPS-treated tricultures did not induce MN cell loss, indicating that this neurotoxic effect is mediated by mechanisms dependent on cell-cell contact (Figure 6D).

**Figure 6.**
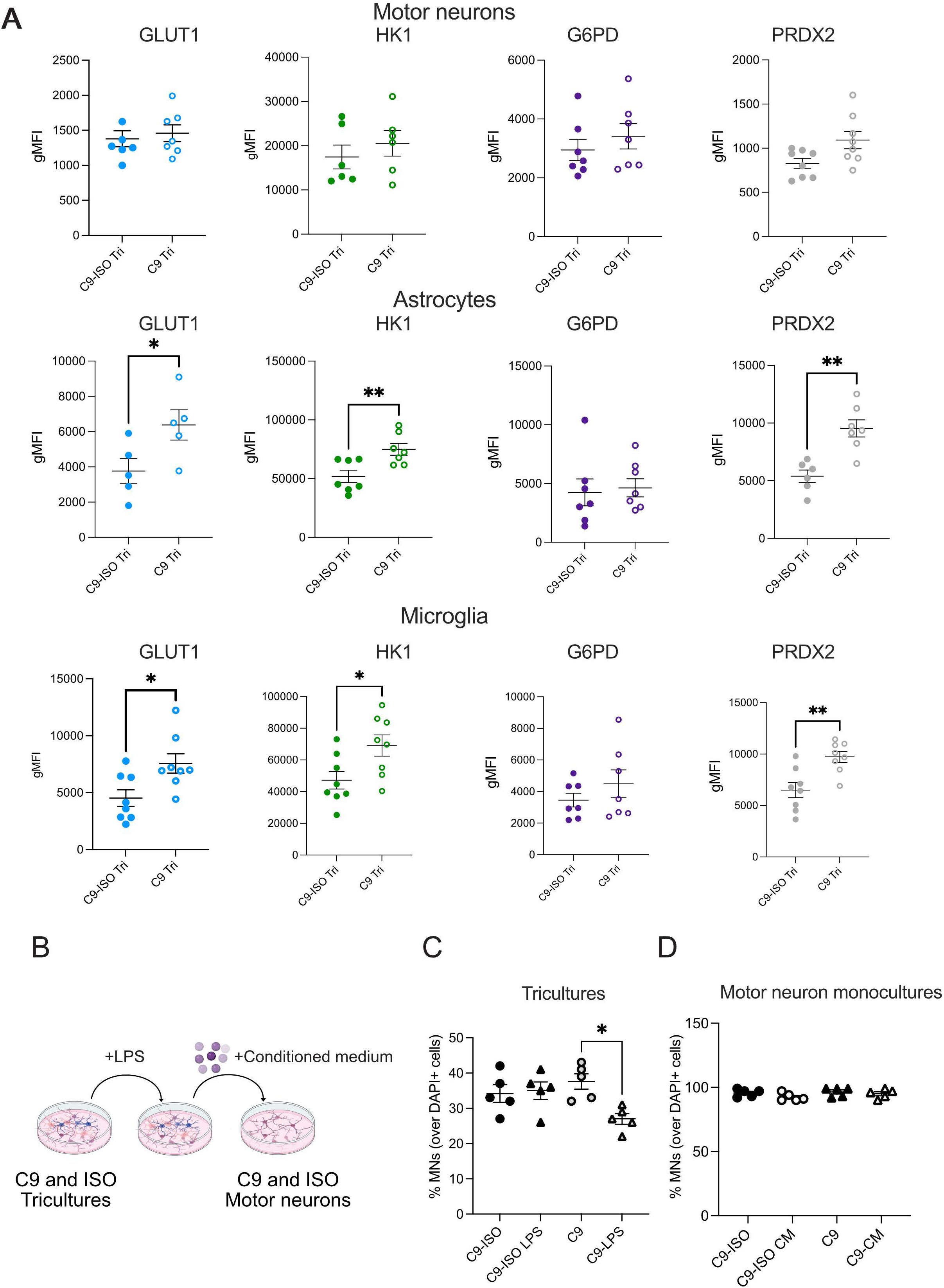
C9orf72 microglia drive astrocytic metabolic changes and contribute to motor neuron loss under inflammatory conditions. **(A)** Met-Flow analysis of metabolic markers in isogenic iPSCs under basal conditions. The metabolic markers GLUT1, HK1, G6PD, and PRDX2 were assessed in MNs, astrocytes, and microglia using Met-Flow. The data are presented as the geometric mean fluorescence intensity (gMFI) for each marker, comparing C9orf72-expanded tricultures (C9) with isogenic controls (C9 ISO Tri) upon LPS treatment. Mean ± SEM; unpaired two-tailed t test; *P < 0.05 and **P < 0.01; n=6-8 independent experiments. **(B)** Schematic of the experimental workflow. C9orf72 and isogenic tricultures were either left untreated or treated with 100 ng/mL LPS for 24 hours. MN monocultures were then exposed to conditioned medium from the corresponding LPS-treated tricultures. **(C)** Quantification of the number of MNs in tricultures with and without 100 ng/mL LPS. **(D)** Quantification of the number of MNs in monocultures with and without treatment with LPS-treated triculture-conditioned media. In C) and D), neuronal numbers were quantified by counting the number of TUBB3-positive neurons in 10 randomly selected fields per sample, normalized to DAPI-positive cells. Mean ± SEM; one-way ANOVA with Bonferroni post hoc correction; *P < 0.05; n=5 independent experiments.

## Discussion

This study highlights the metabolic vulnerabilities associated with *C9orf72* hexanucleotide repeat expansion in ALS, emphasizing the cell type-specific effects on MNs, astrocytes and microglia. Our findings highlight the critical role of intercellular interactions and metabolic regulation in ALS pathogenesis and offer new perspectives for potential therapeutic strategies. Met-Flow, a single-cell metabolic analysis tool, has emerged as an invaluable technique for dissecting the complex interplay of cell type-specific metabolic vulnerabilities. By providing insights at the single-cell level, Met-Flow has helped bridge gaps in the understanding of the metabolic crosstalk between glia and neurons, highlighting potential therapeutic targets aimed at restoring cellular homeostasis and mitigating disease progression.

A key observation was the differential metabolic impact of the *C9orf72* mutation across cell types. While MNs exhibited impaired mitochondrial respiration and reduced ATP production, which are indicative of energy metabolism deficits, microglia exhibited increased glycolytic activity and oxidative stress. Those results suggest that C9orf72 loss of function induces distinct metabolic reprogramming in each cell type. Importantly, the pronounced upregulation of GLUT1 and PRDX2 expression in microglia, both at basal levels and under inflammatory conditions, highlights the heightened metabolic and oxidative stress responses in these cells. The observed effects in microglia are particularly notable given the high expression of C9orf72 in myeloid cells, which are essential for antigen presentation and immune regulation. Consistent with those findings, previous studies have shown that C9orf72 loss leads to hyperactive autoimmune responses and excessive cytokine release in bone marrow-derived macrophages [11, 22, 23]. Interestingly, our results suggest that the C9orf72 mutation does not uniformly affect metabolism in all cell types but instead triggers context-dependent metabolic responses that vary depending on both the cell type and the environmental conditions. To investigate those dynamics, we developed a novel triculture system comprising MNs, astrocytes and microglia; this model closely mimics cellular interactions within the CNS, allowing us to dissect the intricate crosstalk among these cell types. By promoting cell‒cell interactions, the triculture system provides a powerful platform for studying the specific effects of the *C9orf72* mutation on cellular metabolism in a more physiologically relevant context. Our results indicated that the triculture system not only enhanced the metabolic activity of MNs compared with monocultures but also highlighted how the mutation differentially affects the metabolic profiles of each cell type. For example, while MNs in monocultures presented significant mitochondrial deficits, they presented increased metabolic activity in the triculture system, suggesting a potential compensatory effect mediated by neuronal‒glial interactions. Similarly, astrocytes in the triculture system exhibited increased glucose metabolism, as evidenced by the upregulation of GLUT1 expression, suggesting that the multicellular environment modulates their metabolic state. Interestingly, recent data show that partial reduction of astrocytic GLUT1 paradoxically improves glucose utilization in the brain, leading to increased neuronal activity without inducing metabolic stress [24]. Triculture systems may therefore enhance this beneficial astrocytic metabolic feature.

These findings support the growing concept that neuronal–glial crosstalk plays a critical role in shaping the metabolic and functional properties of CNS cells in both health and disease [20]. Importantly, this approach highlights the need to study neurodegenerative diseases in complex, multicellular environments to better understand the interplay between genetic and metabolic factors in disease progression. The metabolic changes observed in microglia are particularly noteworthy in the context of ALS-associated neuroinflammation. The significant upregulation of the expression of glycolytic enzymes (e.g., GLUT1 and HK1) and oxidative stress markers (e.g., PRDX2) in C9orf72 microglia upon LPS stimulation suggests that these cells undergo metabolic reprogramming in response to inflammatory signals. This metabolic shift likely exacerbates oxidative stress and contributes to the neurotoxic environment observed in ALS. Moreover, the ability of activated C9orf72 microglia to drive astrocytic metabolic changes underscores the potential role of microglia‒astrocyte crosstalk in disease progression. This interaction may facilitate the transition of astrocytes from a neuroprotective phenotype to a neurotoxic phenotype, as previously reported in ALS models [21]. Importantly, the increased metabolic activity in astrocytes and microglia appears to precede MN loss, suggesting that these glial cells may play a causative role in neurodegeneration. Our triculture system demonstrated that the presence of C9orf72 microglia and astrocytes significantly increased the vulnerability of MNs to inflammatory stimuli. Interestingly, these effects were mediated by cell-cell contact mechanisms. Our findings support the non-cell-autonomous hypothesis of ALS, in which glial cells actively contribute to MN degeneration through metabolic and inflammatory mechanisms [1]. Furthermore, our results highlight the critical role of neuron–glia interactions in the initiation of neurodegenerative disease processes.

While our study provides valuable insights, it also raises several questions. First, the mechanisms by which the C9orf72 mutation drives metabolic reprogramming in microglia and astrocytes remain incompletely understood. Given the known role of C9orf72 in mitochondrial function [25, 26], perturbations in these pathways may contribute to the observed metabolic changes. However, further studies are needed to elucidate the molecular links between C9orf72 dysfunction and immune metabolic reprogramming. Second, our findings highlight the importance of the cellular microenvironment in modulating the effects of the C9orf72 mutation. The triculture system used in this study provides a valuable platform for studying these interactions, but additional models that incorporate more complex features of the CNS, such as blood‒brain barrier components or peripheral immune cells, may offer further insights. Finally, our study focused on basal and LPS-induced inflammatory conditions. Future investigations should explore how other pathological stimuli, such as oxidative stress or protein aggregation, influence the metabolic and functional properties of C9orf72 mutant cells. Such studies may help identify common pathways that could be targeted therapeutically across different ALS subtypes.

## Conclusions

In conclusion, our findings reveal a critical role for C9orf72 in modulating the metabolic and inflammatory states of MNs, astrocytes, and microglia. The cell type-specific metabolic changes observed in this study highlight potential therapeutic targets for ALS, particularly those aimed at restoring metabolic homeostasis and mitigating oxidative stress in microglial cells. By advancing our understanding of the interplay among metabolism, inflammation, and intercellular communication in ALS, this work provides a foundation for the development of novel strategies to slow or halt disease progression.

## Methods

### Induced pluripotent stem cell (iPSC) culture

The following human C9orf72 iPSC lines and corresponding isogenic controls were used in this study: CS52iALS-C9nxx/CS52iALS-C9n6.ISOxx and CS29iALS-C9nxx/CS29iALS-C9n1.ISOxx (obtained from Cedars-Sinai, Los Angeles, CA, USA) and BS6/BS62H9 iPSCs (Selvaraj et al., 2018). Human iPSCs were maintained on Matrigel (Corning) in mTeSR Plus medium (STEMCELL Technologies) supplemented with penicillin‒streptomycin. Cultures were routinely tested for mycoplasma contamination using a Venor® GeM Classic detection kit (11-1050, Minerva Biolabs GmbH).

### Differentiation of human iPSCs into motor neurons

MN differentiation was performed in accordance with a previously established protocol [27]. Briefly, confluent iPSC cultures were detached from Matrigel-coated plates and dissociated into single cells using Accutase (Stem Cell Technologies) at 37°C for ∼5 min. The resulting single cells were seeded in suspension for differentiation at a density of 1 × 10⁶ cells/mL in complete mTeSR media supplemented with 10 ng/mL FGF2 (PeproTech) and 10 μM ROCK inhibitor Y-27632 (Selleck Chemicals) on Matrigel™-coated 6-well plates. At 48 h postseeding (differentiation day 0), mTeSR medium was replaced with N2B27 MN differentiation medium, a 1:1 mixture of DMEM/F12 (Fisher Scientific) and Neurobasal medium (Thermo Scientific) supplemented with 1% N2, 2% B27, 1% Pen-Strep, 1% GlutaMAX, 0.1% β-mercaptoethanol (βME, Life Technologies), and 20 μM ascorbic acid (Sigma Aldrich). On days 0-1, the medium was supplemented with 10 μM SB-431542 (Selleck Chemicals), 100 nM LDN-193189 (Axon MedChem), and 3 μM CHIR-99021 (Sigma Aldrich). On days 2-4, the medium was supplemented with 10 μM SB-431542, 100 nM LDN-193189, 3 μM CHIR-99021, 1 μM retinoic acid (MedChem Express), and 1 μM smoothened agonist (Sigma Aldrich). On day 5, the medium was supplemented with 1 μM retinoic acid and 1 μM smoothened agonist. On day 7, the medium was supplemented with 20 ng/mL brain-derived neurotrophic factor (BDNF, Peprotech). On day 9, the medium was supplemented with 10 μM DAPT (Selleck Chemicals) in addition to the aforementioned factors. On days 11-13, the medium was supplemented with 20 ng/mL glial-derived neurotrophic factor (GDNF, Peprotech). On day 15, embryoid bodies (EBs) were harvested, washed with 1X phosphate-buffered saline (PBS, VWR) (without calcium and magnesium) and dissociated with 0.25% trypsin-EDTA (Life Technologies) and 50 μg/mL DNase I (Worthington Biochemical) for 5 min at 37°C with gentle shaking. Trypsin activity was quenched with fetal bovine serum (FBS; Sigma Aldrich), and the cells were subsequently centrifuged at 400 × g for 5 min. The cell pellet was resuspended in dissociation buffer (5% FBS, 25 mM glucose, and 1% GlutaMAX in PBS) and mechanically triturated with a P1000 pipette. Dissociated single cells were centrifuged (400 × g, 5 min) and resuspended in complete MN medium (Neurobasal medium containing 1% N2, 2% B27, 1% Pen-Strep, 1% GlutaMAX, 1% nonessential amino acids, 0. 1% βME, 20 μM ascorbic acid, 20 ng/mL BDNF, GDNF and CNTF (ciliary neurotrophic factor, Sigma‒Aldrich) and 10 μM UFDU [1:1 uridine (Sigma Aldrich):fluorodeoxyuridine (Sigma Aldrich)]. The cell suspension was filtered through a 40 μm cell strainer, and the cells were counted using an automated cell counter with a 1:1 dilution of trypan blue and plated at the desired density onto tissue culture plates precoated with 25 μg/mL polyornithine (Sigma Aldrich), 5 μg/mL mouse laminin (Life Technologies) and 10 μg/mL fibronectin (Sigma‒Aldrich).

### Differentiation of human iPSCs into microglia

iPSCs were differentiated into microglia following an established protocol with minor modifications [28]. EBs were formed using AggreWell™800 plates (STEMCELL Technologies) and cultured in mTeSR1 (STEMCELL Technologies) supplemented with 50 ng/mL bone morphogenetic protein 4 (BMP4, ImmunoTools), 50 ng/mL vascular endothelial growth factor (VEGF, ImmunoTools) and 20 ng/mL stem cell factor (SCF, ImmunoTools) for 4 days, with 75% of the medium changed daily. On day 4, the EBs were harvested and transferred to 6-well cell culture plates (12-16 EBs/well) in X-VIVO 15 (Lonza) supplemented with 25 ng/mL IL-3 (ImmunoTools), 100 ng/mL macrophage colony stimulating factor (M-CSF, ImmunoTools), 2 mM GlutaMAX (Thermo Fisher), 1% P/S (Merck Millipore) and 0.055 mM β-mercaptoethanol (Sigma‒Aldrich), with weekly medium changes. After 3–4 weeks, floating cells were collected and seeded at 100,000 cells/cm² on Matrigel-coated plates in Advanced DMEM/F-12 (Life Technologies) supplemented with N2 (Thermo Fisher), GlutaMAX, P/S, β-mercaptoethanol, 100 ng/mL M-CSF, 100 ng/mL IL-34 (PeproTech) and 10 ng/mL GM-CSF (ImmunoTools), with the medium changed twice weekly.

### Differentiation of human iPSCs into astrocytes

Human iPSCs were differentiated into astrocytes in accordance with the protocol described by Tcw et al. [29]. To detach iPSC colonies, the cells were treated with 1 mg/ml collagenase (Thermo Fisher Scientific) for 1 h. After collagenase treatment, the cells were suspended in EB medium and transferred to untreated polystyrene plates for 7 days, after which the medium was changed daily. The EB medium consisted of DMEM/F12 (Gibco, 11330032), 20% knockout serum replacement (Gibco, 10828028), 1X GlutaMAX (Gibco, 35050061), 1X MEM nonessential amino acids (NEAAs, Gibco, 11140050), 100 μM β-mercaptoethanol (Gibco, 21985023), 2 μM dorsomorphin (Tocris, 3093) and 2 μM A-83 (Tocris, 692). After 7 days, the EB medium was replaced with neural induction medium consisting of DMEM/F12, 1X N2 supplement (Gibco, 17502048), B27 supplement, 1X NEAAs, 1X GlutaMAX, 2 μg/ml heparin (Sigma) and 2 μM cyclopamine (Tocris, 1623). On day 7, floating EBs were transferred to Matrigel-coated six-well plates to form neural tube-like rosettes, which were maintained for 15 days, with the medium changed every other day. On day 22, the rosettes were mechanically picked and transferred to low-attachment plates (Corning) in neural induction medium supplemented with B27. Neural precursor spheres were differentiated into astrocytes as described by Tcw et al. with minor modifications. Spheres were dissociated with Accutase for 10 min at 37°C and seeded at 15,000 cells/cm² on Matrigel-coated plates in astrocyte medium (ScienCell: 1801, containing astrocyte medium (1801-b), 2% fetal bovine serum (0010), astrocyte growth supplement (1852) and 10 U/ml penicillin/streptomycin solution). Starting on day 2, the cells were fed every 48 h for 20–30 days. Astrocytes were split at 90–95% confluence (approximately every 6–7 days) and reseeded as single cells in astrocyte medium at the same initial seeding density (15,000 cells/cm²).

### Determination of ATP levels

The level of intracellular ATP was determined using an ATP determination kit (A22066; Thermo Fisher Scientific, Waltham, MA, USA 02451) in according with the manufacturer’s instructions. The amount of ATP in each sample was calculated from standard curves and normalized to the total protein concentration.

### Seahorse XFp metabolic flux analysis

The oxygen consumption rate (OCR) and extracellular acidification rate (ECAR) were measured using an XFp Extracellular Flux Analyzer (Agilent Technologies). iPSC-derived MNs, astrocytes, and microglia were cultured on XFp Cell Culture Miniplates (Agilent Technologies) at a density of 100,000 MNs/well for 10 days, 25,000 astrocytes/well for 5 days, or 80,000 microglia/well for 7 days. The OCR and ECAR were measured in freshly prepared medium consisting of phenol-free DMEM (pH 7.4) supplemented with 1 mM GlutaMAX™. Oxygen consumption was analyzed after the sequential injection of oligomycin, carbonyl cyanide m-chlorophenylhydrazone (CCCP), and antimycin A/rotenone. Glycolytic function was evaluated after the sequential injection of glucose, oligomycin, and 50 mM 2-deoxy-D-glucose (all from Sigma‒Aldrich); three measurements were obtained for each condition, each lasting 3 min. The values were normalized to the protein concentration as measured by a BCA assay (Pierce, WI, USA).

### Human iPSC tricultures

For the triculture of iPSC-derived astrocytes, MNs and microglia, the differentiation protocols were followed as described above with minor modifications. Astrocytes were differentiated as described and seeded at a 1:3 ratio from 95% confluent 12-well plates into Matrigel™-coated 6-well plates on day 0 and maintained in astrocyte medium with 0.5% FBS for 3 days. On day 3, MN precursors were seeded at 52,000 cells/cm² on preplated astrocytes (day 11 of the MN differentiation protocol). The medium was then changed on day 11 to MN medium supplemented with 4% astrocyte medium without FBS. On day 9, microglial precursor cells were harvested and seeded onto cocultured astrocytes and MNs at 31,000 cells/cm². The medium was replaced in accordance with the MN maturation protocol and supplemented with 4% astrocyte medium without FBS, 100 ng/mL M-CSF, 100 ng/mL IL-34 and 10 ng/mL GM-CSF. Approximately 50–66% of the medium was replaced on days 11 and 13. On day 15, the cells were ready for subsequent experiments.

### Immunocytochemistry and image analysis

Cells were fixed in 4% paraformaldehyde (PFA) in PBS (w/v) for 15 min, washed with PBS, and blocked with PBST (PBS containing 0.1% Triton X-100) supplemented with 10% normal goat serum (NGS) or 10% normal donkey serum (NDS) for 1 h. The cells were then incubated overnight at 4°C with the primary antibody of interest, which was diluted in PBST containing 5% NGS or NDS. After three 10-min washes in 1x PBS, the appropriate species-specific Alexa Fluor 488/568/647-conjugated secondary antibody (Invitrogen) was applied to the cells for 2 h at room temperature, after which they were resuspended in PBST containing 5% serum. Next, the cells were washed once for 10 min in 1x PBS, followed by 10 minutes of DAPI staining (1:10,000) and three additional 10-min washes. The cells were mounted with fluorescent mounting medium (Agilent Technologies). The primary antibodies used were rabbit anti-MNX1 polyclonal antibody (1:500, Sigma‒Aldrich #HPA-071717), rabbit anti-CHAT (1:200, Merck Millipore #AB144P), mouse anti-tubulin β III (1:1000, BioLegend #801202), rabbit anti-tubulin β III (1:1000, BioLegend #903401), mouse anti-PU.1 [7C6B05] (1:500, BioLegend #658002), rabbit anti-IBA1 (WAKO 016-2001), chicken anti-GFAP (1:2000, BioLegend # 829401), and mouse anti-GLAST (ACSA-1) (1:50, Miltenyi Biotech # 130-132-738). For LPS treatment, 100 ng/ml LPS (ultrapure LPS from *E. coli* 0111:B4, InvivoGen) was added to mono- or tricultures 24 h prior to sample collection. Where indicated, conditioned medium was collected from control tricultures and tricultures stimulated for 24 h with 100 ng/ml LPS. The conditioned medium was filtered and then used for MN survival assays. Neuronal counts were quantified by counting the number of TUBB3 neurons in 10 randomly selected fields per sample. Each condition was tested in a minimum of three independent experiments. Images were analyzed with Fiji-ImageJ version 2.3.0/1.53q.

### Flow cytometry

Cells were detached with Accutase™, resuspended in 100 µL of PBS, stained with Zombie NIR™ Fixable Viability Dye, incubated with Fc block (BioLegend) for 15 min, counted and incubated for 30 min in the dark with fluorophore-conjugated antibodies. After washing, the cells were fixed in 4% PFA (15 min, RT, dark) after permeabilization with 0.1% Triton-X and 1% FBS in PBS for 15 min at RT in the dark. The cells were then stained with primary antibodies for intracellular staining in permeabilization/blocking buffer and incubated for 30 min on ice, protected from light, followed by a second incubation for 30 min on ice with secondary antibodies. After final washes, the cells were resuspended in PBS containing 2% FBS and transferred to 5 ml round bottom tubes for flow cytometry. Filter cap tubes were used for MN samples to avoid clogging the instrument. The samples were run on a BD LSRFortessa™ and analyzed using FlowJo™ software. The following antibodies were used for flow cytometry: anti-CD11b-BV785 (clone ICRF44, 1:40; BioLegend, # 301346); anti-CD49f PE/Cyanine7 anti-human/mouse (clone GoH3; 1:50; BioLegend, #313622); CoraLite®488-conjugated anti-peroxiredoxin 2 (Clone 3H6C4; 1:50; ProteinTech, # CL488-60202); DyLight 650 anti-Glut1 (GLUT1/2475) (clone GLUT1/2475; 1:50; Novus Biologicals, #NBP2-75786C); Immunotag™ AF405 anti-G6PD monoclonal antibody (1:50; Immunotag™; G-Biosciences; #ITM0292); and Immunotag™ AF594 anti-HXK I monoclonal antibody (1:50; G-Biosciences; # ITM0348). For the Met-Flow experiments, MNs were labeled with 1 µM CellTrace™ Yellow dye (Thermo Fisher Scientific, #C34573) prior to being seeded onto precoated astrocytes. Precursors were detached using Accutase™, counted, and stained in serum-free conditions with 1× PBS for 20 min at 37°C with 5% CO₂. The staining process was halted by adding 20% FBS for 5 min. The cells were subsequently centrifuged and resuspended in MN medium supplemented with 4% astrocyte medium. High-dimensional analysis was performed using Fast Fourier Transform-accelerated interpolation-based t-distributed Stochastic Neighbour Embedding (FitSNE) in FlowJo (BD, version 10.6.1). FitSNE was applied to a downsampled dataset of 10,000-20,000 cells per sample. The histograms adjacent to the FitSNE plots show the number of fluorescence channels on the y-axis and the biexponential fluorescence intensity of each marker on the x-axis.

### Images

Images were created with BioRender.com.

### Statistics

Statistical differences among groups were evaluated using GraphPad Prism Version 10.2.1. All experiments were performed at least in triplicate, and the quantitative data are presented as means ± SEMs. Sample sizes were chosen on the basis of previous studies. All the samples were included in the analysis. Blinding was not performed in this study. Differences among groups were assessed using unpaired two-tailed Student’s *t* test or ANOVA for multiple comparisons, as indicated in the figure legends.

## Acknowledgements

We thank the FACS Core Facility Location Berg, Eberhard Karls University, Tübingen, Germany.

## Authors contributions

M.D. and S.C. conceived the project. M.D., M.M., I.H., and F.P. designed the experiments and interpreted the results. M.M., I.H., F.P., CW, M.R., L.O. performed and analyzed the data. M.M. drafted the first draft of the figures and manuscript with input from the other authors. M.M. and M.D. wrote the paper. All authors read and approved the final version of the manuscript.

## Availability of data and materials

All data are available in the main text, or in the Supplementary Tables and Supplementary Figures. The raw data that support the findings of this study are available from the corresponding author upon request.

## Abbreviations

ALS: amyotrophic lateral sclerosis
ANOVA: Analysis of variance
CNS: Central nervous system
FTD: Frontotemporal dementia
GLUT1: Glucose transporter 1
G6PD: Glucose 6-phosphate dehydrogenase
HK: Hexokinase-1
iPSCs: Induced pluripotent stem cells
Met-Flow: Metabolic flow cytometry
MN: Motor neurons
OCR: Oxygen consumption rate
PRDX2: Peroxiredoxin 2

## Ethics approval and consent to participate

Not applicable.

## Consent for publication

Not applicable.

