## Supplementary Data for "C9orf72 Repeat Expansion Induces Metabolic Dysfunction in Human iPSC-Derived Microglia and Modulates Glial–Neuronal Crosstalk"

### Supplementary figure legends

**Supplementary Figure 1. ATP concentration and ECAR measurements in motor neurons, astrocytes, and microglia. (A)** ATP concentration (relative to that of the control) measured in MNs, astrocytes, and microglia. Each panel compares C9-1, C9-2, and C9-3 to their corresponding isogenic controls (C9-ISO). The amount of ATP in each sample was calculated from standard curves and normalized to the total protein concentration. Mean ± SEM; unpaired two-tailed t test; ** $P<0.01$, * $P<0.05$; $n=5$ independent experiments. **(B)** Extracellular acidification rate (ECAR) measurements in MNs, astrocytes, and microglia, indicating glycolytic flux over time (in minutes). Comparisons are made between C9-1, C9-2, and C9-3 and their corresponding ISO controls (C9-ISO). Two-way ANOVA with the Bonferroni post hoc correction, * $p<0.05$, $n=5$ independent experiments.

**Supplementary Figure 2. High-resolution Met-Flow single-cell metabolic analysis of C9orf72 microglia. (A)** Representative t-SNE plots depicting the expression levels of PRDX2 in C9orf72 and isogenic control microglia under basal conditions. **(B)** Representative t-SNE plots depicting GLUT1 and G6PD expression in C9orf72 microglia under basal conditions and after 100 ng/mL LPS stimulation. The color gradient from green to red represents the intensity of expression, with green indicating lower expression and red indicating higher expression.

**Supplementary Figure 3. Met-Flow metabolic analysis of iPSC-derived neuronal-glial tricultures. (A)** Gating strategy used to perform Met-Flow metabolic analysis of iPSC-derived astrocytes, MNs and microglia in isogenic tricultures. Representative scatter plots illustrate the sequential gating approach in C9orf72 and isogenic control iPSC neuronal glial tricultures (isogenic and mutant, top and bottom rows, respectively). Cells were first gated for viability and singlets, followed by cell type identification: microglia (CD11B+), astrocytes (CD49F+) and MNs (CellTrace+). MNs were further distinguished by the absence of both CD11b and CD49f expression. The scatter plots show the forward scatter (FSC-A) and CellTrace parameters, allowing a clear separation of microglia, astrocytes and MNs. **(B)** t-SNE plots of phenotypic markers showing the spatial distribution of astrocytes (orange), microglia (blue) and MNs (green) on the basis of CD49f, CD11b and CellTrace markers, confirming cell identity and purity.

**Supplementary Figure 4. Metabolic assessment of iPSC-derived neuronal and glial cells in monocultures versus tricultures with C9orf72 expansion under basal conditions using Met-Flow.**

Quantitative analysis of the metabolic markers GLUT1, HK1, GSPD and PRDX2 in MNs, astrocytes and microglia derived from the C9orf72 isogenic (C9-ISO) and expanded C9orf72 (C9) iPSC neuronal–glial monocultures and tricultures under basal conditions. The data are expressed as the geometric mean fluorescence intensity (gMFI). Mean ± SEM; unpaired two-tailed t test; ***P < 0.001, **P < 0.01, and *P < 0.05; n = 6-8 independent experiments.

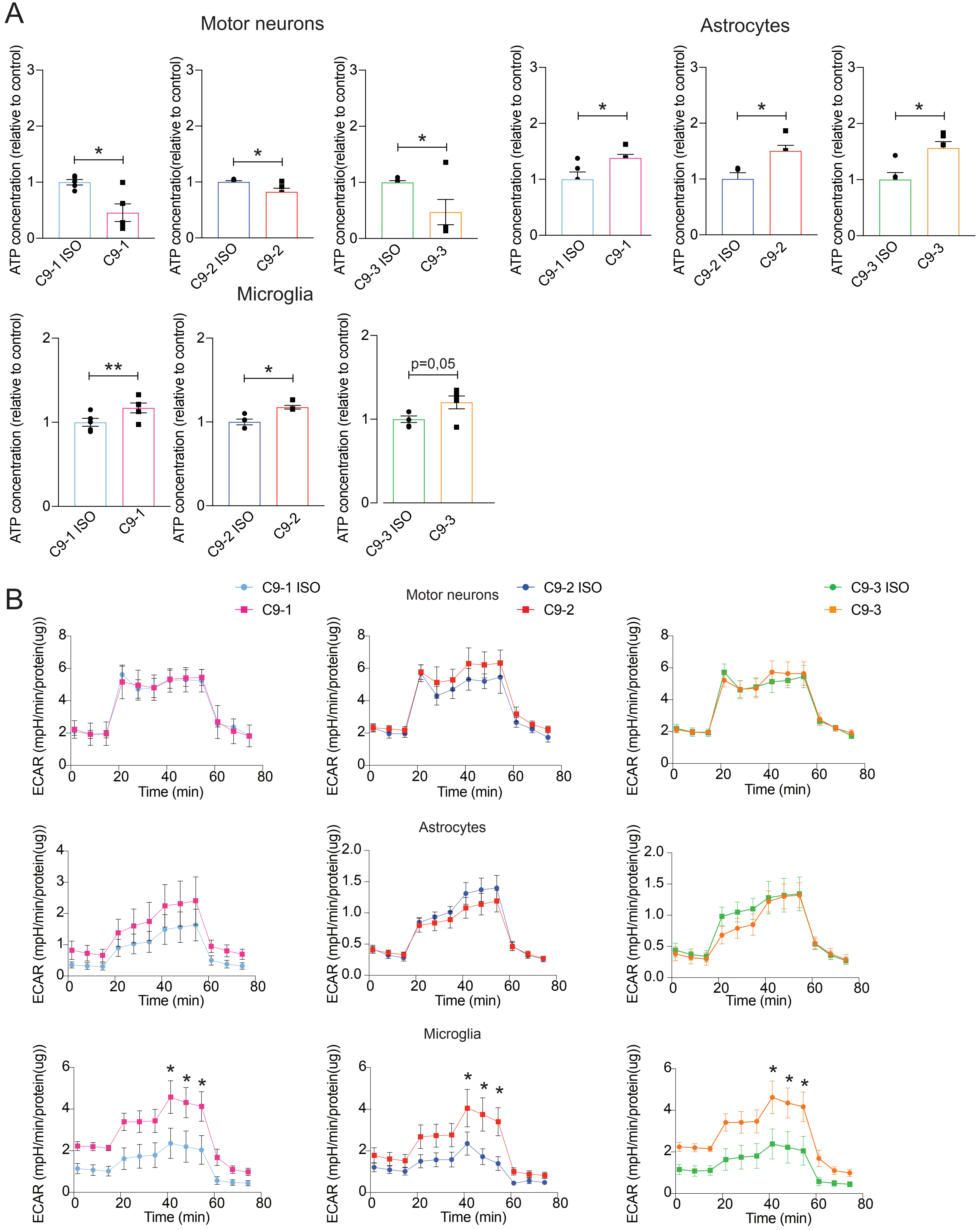


Supplementary Figure 1

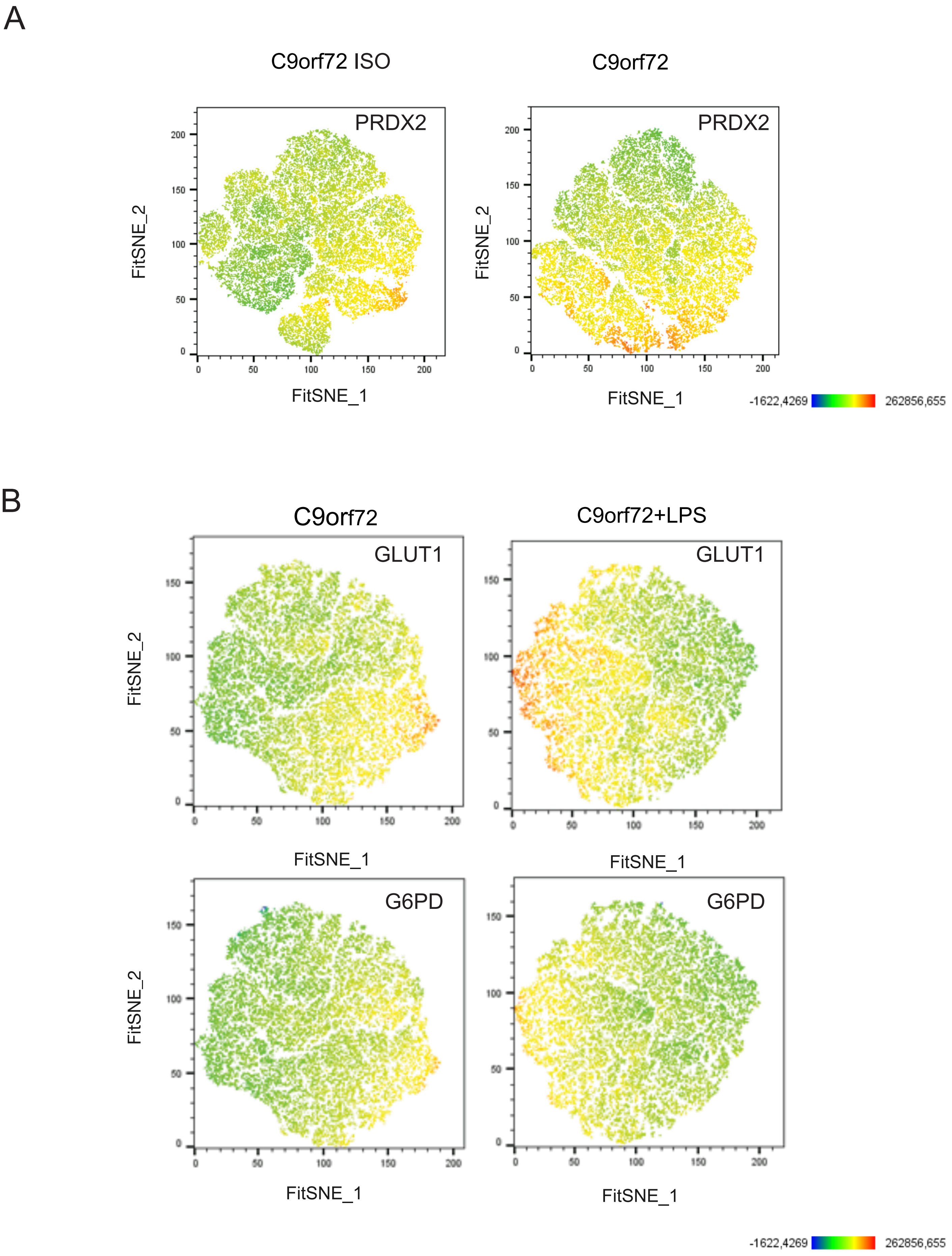


Supplementary Figure 2

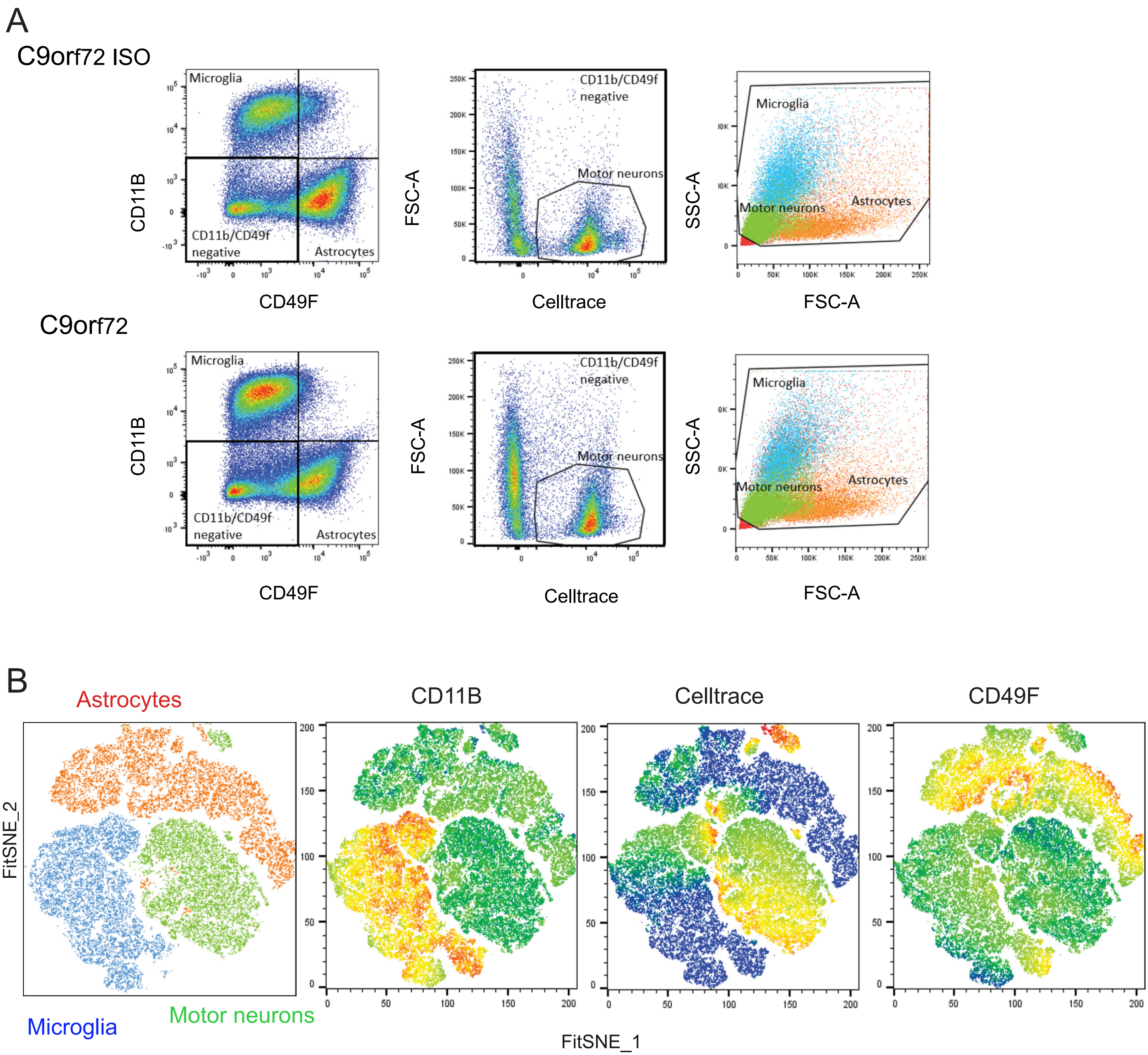


Supplementary Figure 3

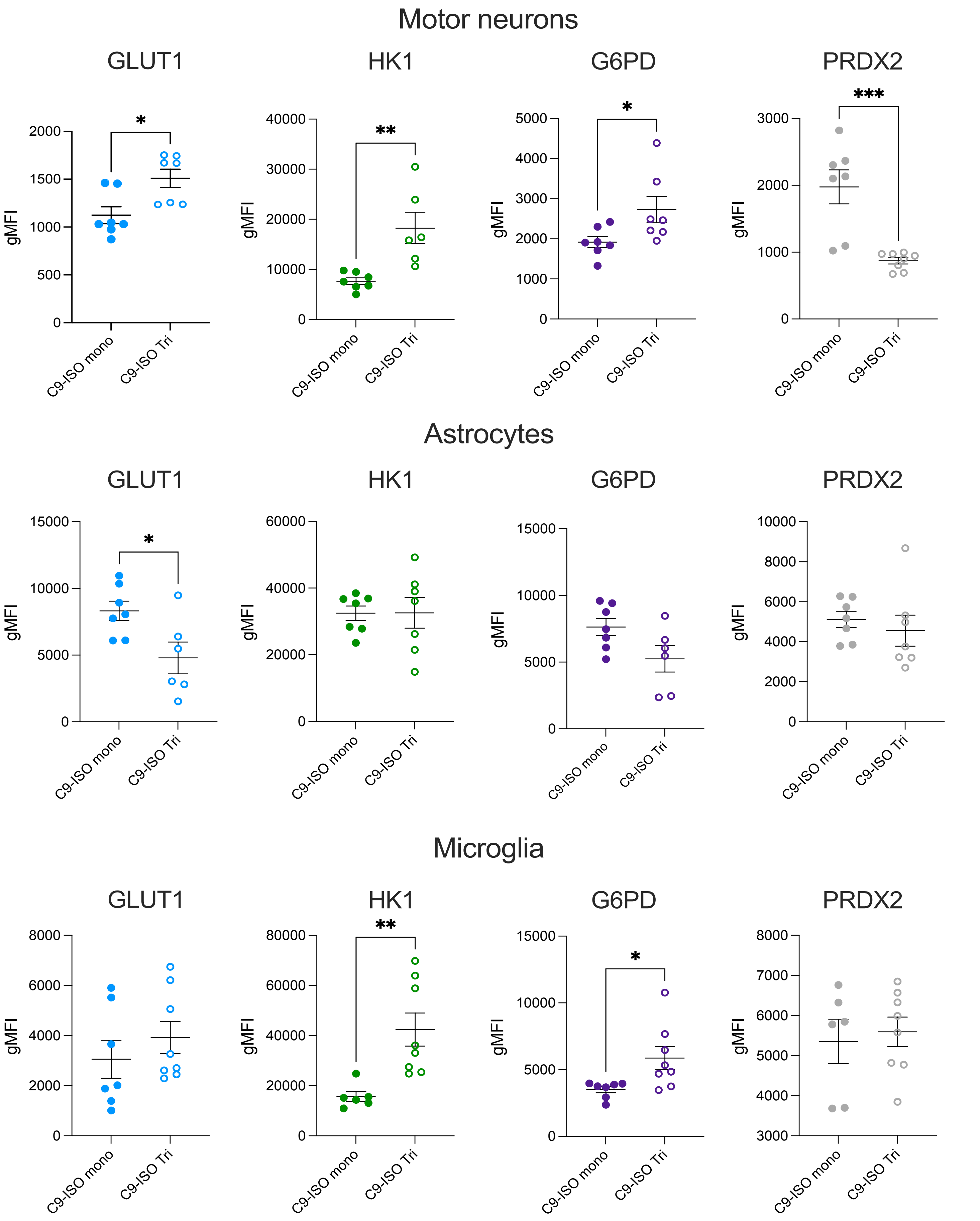


Supplementary Figure 4
